# Combining CRISPR/Cas mediated terminal resolution with a novel genetic workflow to achieve high diversity adenoviral libraries

**DOI:** 10.1101/2023.11.16.566979

**Authors:** Julian Fischer, Ariana Fedotova, Lena Jaki, Erwan Sallard, Anja Erhardt, Jonas Fuchs, Zsolt Ruzsics

## Abstract

While recombinant Adenoviruses (rAds) are widely used in both laboratory and medical gene transfer, library-based applications using this vector platform are not readily available.

Recently, we developed a new method, the CRISPR/Cas9 mediated in vivo terminal resolution (CTR) aiding high efficiency rescue of rAds from recombinant DNA. Here we report on a genetic workflow that allows construction of BAC-based rAd-libraries employing the efficiency of CTR.

We utilized frequent, pre-existing genomic sequences to allow insertion of a selection marker, complementing two selected target sites into novel endonuclease recognition sites. In a second step, this selection marker is replaced with a transgene or mutation of interest via Gibson assembly. Our approach does not cause unwanted genomic off-target mutations while providing substantial flexibility for the site and nature of the genetic modification.

This new genetic workflow, which we termed half-site directed fragment replacement (HFR) allows introduction of >10^6^ unique modifications into rAd encoding BACs using laboratory scale methodology. To demonstrate the power of HFR, we rescued barcoded viral vector libraries yielding a diversity of ∼2.5×10^4^ modified rAd per cm^2^ of transfected cell culture.

**GRAPHICAL ABSTRACT:** 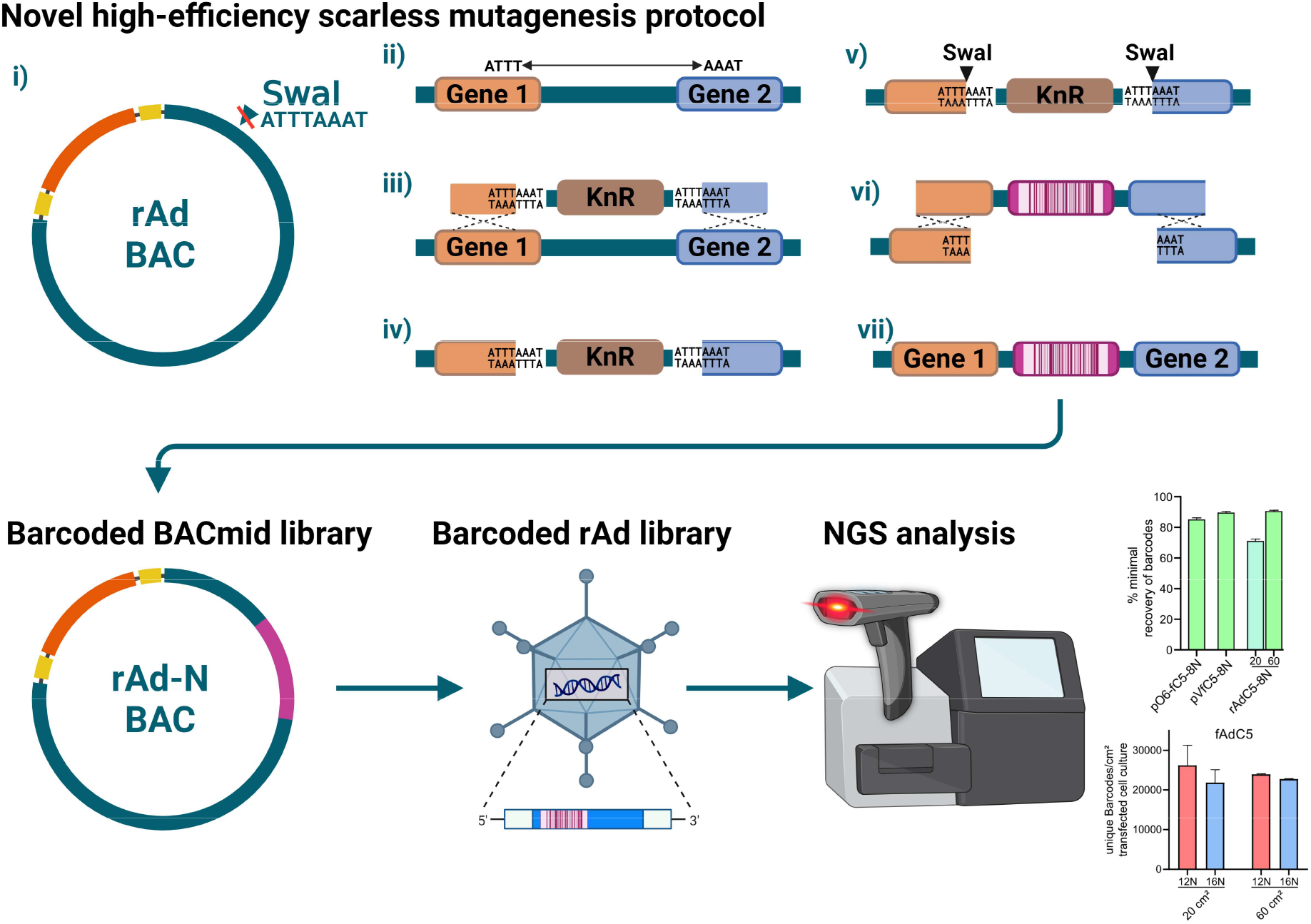

## INTRODUCTION

Worldwide, 14 different viral gene therapies are currently approved for clinical use in humans (1, 2). Half of these therapies are based on either human adenoviruses (HAdV) or chimpanzee adenovirus and there are numerous clinical trials and even more research applications with recombinant adenovirus (rAd)-based vectors. Due to their large capacity for transgenes, simple cloning- and production-principle and ability to mediate efficient gene transfer both in vitro and in vivo, rAds are an attractive platform for gene transfer applications (3–5).

During recent years, recombinant adeno-associated viruses (rAAV) vector technology allowed for randomized recombination of viral serotypes, generating a huge amount of novel viral capsids (6, 7). These feature libraries paved the way to screen for variants with novel tropism, allowing for the targeting of specific cells while minimizing unwanted off-target effects (8–10). This technological advantage promoted the rAAV-platform to the method-of-choice among the currently ongoing gene therapy clinical trials (1). For rAds, the method of choice for finding novel vector variants remains rational design (5, 11, 12). Despite being successful in some cases, rational design is limited by low throughput and, in the case of vector retargeting, by potentially unknown targeting sites.

In order to allow library reconstitution, an effective method is required to achieve high yield virus preparations carrying diversified genetic material. Recently, we developed a new method to rescue rAd that performs comparable to rAAV or recombinant lentivirus rescue. This technology is based on CRISPR/Cas9-mediated terminal resolution (CTR) and allows an efficient rAd rescue in different cell types (13). Notably, AAV- or lentivirus vector genomes can be easily modified and maintained in small high-copy plasmids in E. coli, simplifying library construction. In contrast, modification of large rAd genomes rarely reaches efficiencies that would permit library applications.

Here, we describe a novel method to reliably modify bacmids at any given position. Utilizing CTR, we demonstrate the rescue of infectious and barcoded rAd populations that surpass the variability of currently established rAd library methods.

## MATERIAL AND METHODS

Tables for diversified oligonucleotides, primers and plasmids used in this study can be found in supplementary data S1-3.

### Cell and viruses

293A (Thermo Fisher Scientific, Waltham, MA, USA, Invitrogen #R70507) and A549 (ATCC, Manassas, VA, USA, CCL-185) cells were cultured using Dulbecco’s modified Eagle’s medium (DMEM; Anprotec, Bruckberg, Germany, #AC-LM-0014) supplemented with 10% fetal calf serum (FCS; PAN-Biotech, Aidenbach, Germany, #P30-3306) and penicillin–streptomycin (100 U/ml; Thermo Fisher Scientific, Waltham, MA, USA, gibco #15140-122). The wild type HAdV-C5 was cultured from inoculum provided by Albert Heim (German Adenovirus Reference Laboratory, Hannover). HH-Ad5-VI-wt (14) was cultured from an inoculum provided by Harald Wodrich (Microbiologie Fondamentale et Pathogénicité, Bordeaux).

### Bacterial Strains

| Strain | Features | Product ID |
| --- | --- | --- |
| *E. coli* NEB10beta | $\Delta$(ara-leu) 7697 araD139 fhuA ΔlacX74 galK16 galE15 e14- Φ80dlacZΔM15 recA1 relA1 endA1 nupG rpsL (StrR) rph spoT1 $\Delta$(mrr-hsdRMS-mcrBC) | NEB C3020K |
| *E. coli* Pir1 | F- Δlac169 rpoS(Am) robA1 creC510 hsdR514 endA1 recA1 uidA(ΔMluI)::pir-116 | Invitrogen™ C101010 |

### Construction of recombinant bacmids and transfer plasmids used for library assembly

First, CTR-compatible bacmids for replication competent and first generation recombinant human adenovirus C5 genomes were generated as previously described (13). To construct the bacmid carrying the wt genome, (pBWH-C5), we fused PCR amplified vector fragments based on pKSB2 (15) backbone using primer pairs BWHC05for/GHBrev and GHBfor/BWHC05rev with viral genomic DNA isolated from infected cells. We used the same approach to clone an E3-deleted genome using viral DNA isolated from cells infected with HH-Ad5-VI-wt, which is a replication competent HAdV-C5 strain, which carries an E3 deletion (14). Finished constructs were verified using restriction fragment length polymorphism (RFLP) analysis, sequencing and checked for viability using CTR.

For completing the half sites to new recognition sequences of restriction endonuclease SwaI, which we utilized in our new genetic workflow, we used recombineering as described by Datsenko and Wanner (16). As linear fragments for recombination, gene cassettes encoding for kanamycin (Kn) resistance flanked by the completed SwaI sites were amplified via PCR on a pGPS1.1 template. Primers were designed to contain 30 nt 5’-homologous regions to genomic regions targeted for insertion as well as 4 nt sequences completing SwaI restriction sites in addition to their priming sites for the Kn-cassette. For construction of the pIX-modification designated intermediate, primers C5-pIXKan_for and C5-pIXKan_rev, for barcoding library designated intermediate primers fC5-Kan-L_for and fC5-Kan-L_rev were used. Recombination competent bacteria carrying the target genomes were transformed by the linear fragments. Resulting recombinants were selected for both Kn and chloramphenicol (Cm) resistance and analyzed using RFLP. Correct insertion was furthermore confirmed using Sanger sequencing. Verified molecular clones for the pIX tagging were coined pBWH-C5-pIX-Kn. In generating pBWH-fC5-E1Kn for the library construction, we also deleted the E1 region, which resulted in a new bacmid to construct first generation rAd vectors in the next step.

In order to provide high quality linear inserts for efficient assembly into genomic bacmids, we generated an optional transfer vector for use with barcoding libraries, which could release its transgenic insert by restriction endonuclease digest. For this, synthetic dsDNA inserts containing SapI site flanked genomic sequences between the targeted SwaI half-sites including adjacent homologies, which need to be retained in the first generation vectors, a GFP expression cassette as well as a multiple cloning site (MCS) for the insertion of the randomized sequences have been inserted into pO6-A5-GFP (17) resulting in pO5-fC5-GFP. SapI sites allowed the release of the exact fragment that could be used directly for the genomic assembly. Correct construction was confirmed using RFLP and Sanger sequencing.

### Construction of pO6-fC5-GFP-MCS based diversified barcoded shuttle plasmid libraries

To generate a controlled source of barcoded linear fragments first we constructed a barcoded pO6-fC5-GFP-MCS shuttle plasmid library by Gibson assembly (18). Linearized vector fragments were amplified by PCR using primers PCR-LibBB_for and PCR-LibBB_rev. The inserted barcoding sequences were annealed using randomized overlapping single stranded oligonucleotides by adding 3 µl of each oligonucleotide (100 µM) to 500 µl 1x 2.1 NEB-buffer (New England Biolabs, Ipswich, MA, USA, #B6002S) followed by boiling at 100 °C for 3 minutes to finally let them associate during the cooling to room temperature. 2-4 diversified oligonucleotides (see supplementary data S1) were used for each library. General design of oligonucleotides consisted of 30 nt homologous regions on both ends facing to either adjacent oligonucleotides or vector backbone, as well as a 6 nt central diversity cassette (ATNNNNTA). Finally, 100 ng purified linear vector fragments were assembled with 5 µl of annealed oligonucleotide pool using NEBuilder HiFi DNA Assembly (New England Biolabs, Ipswich, MA, USA, #E2621L) according to the manufacturer instructions. The assembly mixes were then dialyzed against deionized water, and competent E. coli Pir1 cells were electrotransformed by 2 µl of the dialyzed assembly mixes. The transformed bacteria were struck out on Kn-containing LB-agar plates and incubated overnight. The entire pool of these bacteria was harvested by adding liquid LB media onto the plate, tapping it lightly and transferring the media to inoculate 100 ml cultures for large scale plasmid DNA preparation of the recombinant transfer plasmid libraries.

The DNA extracted and purified from these pools was then digested using HincII (New England Biolabs, Ipswich, MA, USA, #R0103S) and SapI (New England Biolabs, Ipswich, MA, USA, #R0569S) overnight in order to release transgenic insert and remove vector background. After heat inactivation of the restriction endonucleases (20 min. at 65 °C) these DNA preparations were directly used in the bacmid assemblies as barcoding inserts.

### Construction of genomic bacmid assemblies

The targeted intermediate bacmids pBWH-C5-pIX-Kan or pBWH-fC5-E1Kn were digested by SwaI (New England Biolabs, Ipswich, MA, USA, #R0604S) overnight and heat inactivated at 65 °C after which these preparations were used as vectors in the genomic assembly reactions. The genomic assemblies were set up combining ∼400 ng of bacmid vector preparation and either 15 ng of each PCR derived insert fragments (for pIX modifications) or 100 ng barcoding insert (for the library assemblies) at a final volume of 20µl in the presence of 1x NEBuilder HiFi DNA Assembly Master Mix on ice. The reaction was incubated 60 min at 50 °C and dialyzed against deionized water. Electrocompetent E. coli NEB10beta (New England Biolabs, Ipswich, MA, USA, #C3020K) were transformed by 4 µl of dialyzed bacmid assemblies according to the manufacturer instructions and plated on chloramphenicol (25 µg/ml)-containing LB-agar plates and incubated overnight. For the pIX-modification experiments, single clones were isolated, analyzed by RLFP and finally verified by Sanger sequencing. For the barcoding libraries the entire pool of plated bacteria was harvested by adding liquid LB media onto the plates, tapping it lightly and transferring the media to inoculate 200 ml cultures for large scale low copy plasmid DNA preparation by column purification using NucleoBond Xtra Midi kit (Macherey Nagel, Düren, Germany, #740410.50), analyzed by next generation sequencing and used directly for rAd rescue by CTR.

### Rescue of viral libraries & extraction of viral genomes

For rescuing rAds from the assembled bacmids, Cas9-mediated terminal resolution (CTR) was performed as previously described (13). Shortly, 10 cm^2 cell culture surface of subconfluent 293A cells were co-transfected by 1 µg purified genomic bacmid DNA (gained either from single clones or bacmid library preparations) and 0.5 µg pAR-gRNA-Int5 Cas9-expressing helper plasmid via Lipofectamine 2000 (Thermo Fisher Scientific, Waltham, MA, USA, #11668019). 24 hours later the transfected cells were split either 1:16 into 96-well plates-for isolation of single rescues or completely transferred to a 10 cm dish (for library rescues). Cells were monitored using fluorescent microscopy. For rAd-library applications, the cultured cells were harvested 4 days post transfection. The cells were collected by centrifugation and re-suspended in PBS and frozen using liquid nitrogen. After two freeze-thaw cycles, samples for titration of infectious viruses were saved. The bulk of the lysates was used for viral DNA preparation. Benzonase (Sigma-Aldrich, St. Louis, MO, USA, #E1014-25KU) was added to the lysates to remove both genomic DNA and remaining transfected bacmid. Benzonase was inactivated using 50 mM EDTA and DNA was then extracted using Monarch Genomic DNA purification kit (New England Biolabs, Ipswich, MA, USA, #T3010S).

The single rescues were harvested from single wells containing a single focus of reconstituted virus after full lysis of the identified wells occurred (the first foci appeared 2-3 days after transfections and lysis was observed regularly at 7-8 day after transfection). These primary lysates were expanded to high titer preparations.

### Virological methods

The viral titers were determined by TCID-50 assay using an AAV-based replicon reporter (19). The thermostability of the purified viruses were determined as follows: Viral lysates were incubated at 45 °C for up to 10 min as well as an aliquot at 55 °C for 10 min as described by Vellinga et al. (20). Afterwards, the infectivity of the treated lysates was compared to untreated controls using the TCID-50 assay.

Multistep viral growth analysis was performed using 293A cells which were infected at MOI 0.1 of the respective rAd and incubated for 90 min. Zero-point samples were collected using both culture supernatant and infected cells, remaining samples for later time points were washed with PBS and fresh media was added. Thereafter, supernatants and cells were both collected at time points of up to 72 h post infection, the samples were lysed by 3 cycles of freezing and thawing and total titers were determined by titration.

### Next generation sequencing

For next generation sequencing, NEBNext ULTRA II FS DNA Library Prep (New England Biolabs, Ipswich, MA, USA, #E6177) was used. For diversity analysis on viral bacmid or plasmid DNA, a PCR was performed using primers M13A_for/M13A_rev targeting only N-diversified regions to maximize reads. Indexed pair-end libraries were prepared for Illumina sequencing and normalized and pooled sequencing libraries were denatured with 0.2 M NaOH. Sequencing was performed on an Illumina MiSeq instrument using the 300-cycle MiSeq Reagent Kit v2 (Illumina, San Diego, CA, USA, MS102-2002).

### Data analysis and statistics

NGS data was initially pre-processed and analyzed using a previously published (21) Galaxy (22) workflow. Sequencing adapters were trimmed and low quality basecalls (phred Q < 30) discarded. Reads were then mapped to a reference of the diversified genomic regions and resulting sequences were extracted. Using an in-house R script, single 4-nt diverse regions were extracted and matched to each other, resulting in barcodes 8-16 nt in length, depending on library assembly. Reads not providing a full barcode were discarded. Remaining barcodes were then analyzed further. This was performed to a total of 3 times for each pool. Using lists of unique barcodes from each run for a single sample, total diversity of library pool could be estimated using a heterogeneous capture-recapture model established by Chao (1987)(23). Variant call frequencies of full genome sequencing were cut off at a minimum coverage of 25 reads and showing variants with a frequency >0.1.

### Software

In silico sequence analysis and planning was performed using Geneious Prime (24). NGS data was processed using custom Galaxy (22) and R (25) workflows. Capture/Recapture models were performed using R package Rcapture (26). Sequence logos were generated using R package ggseqlogo (27). Variant frequency plots were generated using VirHEAT (28). Mapping data was generated using QualiMap 2 (29). Graphical Abstract was generated using BioRender.com.

## RESULTS

### Establishment of half-site mediated fragment replacement (HFR) for library efficiency manipulation of rAd-bacmids

Among rAd modification technologies only the red recombineering method allows a free choice of the mutagenesis site but relies on a counter-selection process, making it unreliable in library applications (16). Other methods, which can be performed with higher reliability depend on fixed sites or modification of pre-determined fragments (30–32). This restricts the location of the genetic modification to limited sites either determined by the genomic sequences or inserted artificially, since a unique entry point is required. In both cases it is mandatory to ensure no off-target effects to either adjacent or targeted genomic sequences. We wanted retain the flexibility of red recombineering in respect to choosing the mutagenesis site, so we decided to use existing genomic features which can be found frequently within genomic sequences to provide flexibility of targeting the manipulations. After initial recombination, this workflow would then utilize more reliable methodologies such as Gibson assembly (18) to generate final recombinants. Our aim was to tailor the HAdV-C5 genome for Gibson assembly allowing in vitro manipulation of linear sequences. For this purpose, restriction enzyme recognition sequences which are natively not found within the genomic constructs have to be used. One of the commercially available restriction endonucleases SwaI cannot cut pBWH-C5-delE3 due to the absence of the ATTTAAAT recognition site (Figure 1 A). Instead of artificially inserting the 8 nt site, we analyzed the sequence of the pBWH-C5-delE3 for the frequency of half sites of the SwaI recognition sequence (ATTT and AAAT). Theoretically, one of these sites occurs every 4^4^ or 256 nucleotides. Within the analyzed 40498 bp construct, we found 112 forward facing half sites (ATTT) and 125 backwards facing half sites (AAAT) within the rAd genomic region. The average distance ranged between 2093 nt and 6 nt for the forwards sides and between 2423 nt to 5 nt for backward-sites, averaging at 303 nt and 270 nt, respectively (Figure 1 B).

**Figure 1:**
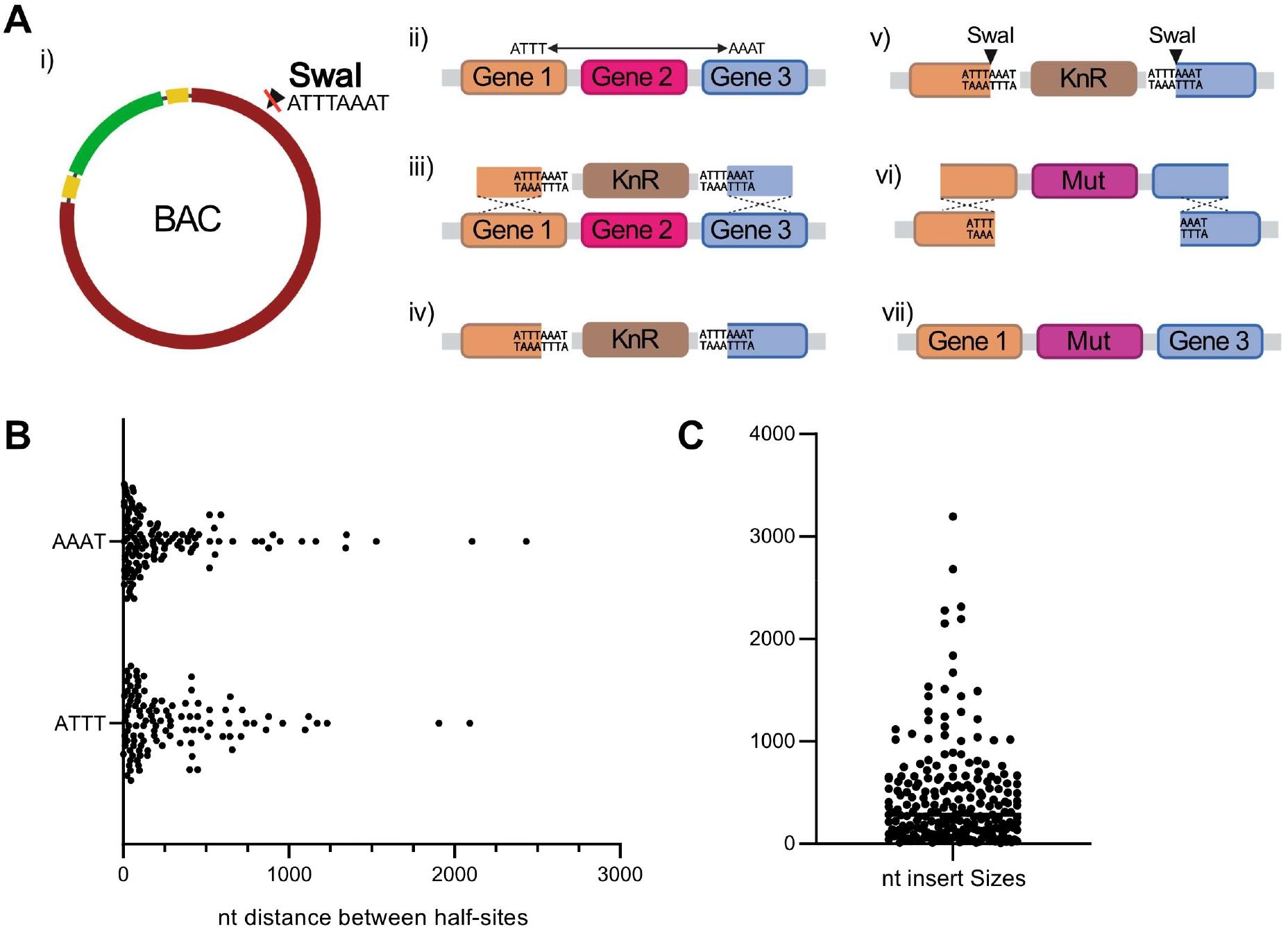
(A) (i) Basic cloning principle requires a bacmid containing viral genome, which is not cut by target enzyme (e.g. SwaI). (ii) Insertion site is determined by selection of target region and offset half sites. (iii) red recombineering is used to replace target region with Kn resistance, (iv) completing proximal SwaI half sites. (v) Intermediate construct is then digested using target enzyme, linearizing plasmid and (vi) allowing insertion of target DNA fragment, resulting in (vii) scarless modified DNA. (B) Distance in nucleotides between half-sites of SwaI in exemplary bacmid containing HAdV-C5delE3 genomic DNA. (C) Minimal required insert size for re-assembly of bacmid (see A-vi)

To create the full SwaI sites based on these half sites one needs to use one forward facing and one reverse facing site in combination as target for red recombineering. Of course, if a SwaI site from two both half sites is generated the central sequence between both sites will be lost and needs to be reinserted together with the modified feature(s) to maintain the integrity of the viral genome. In pBWH-C5-delE3, the required minimal re-insertion fragment sizes ranged between 6 nt and 3191 nt (Figure 1 C). Given these size constraints, the re-insertion fragment can be a synthetic dsDNA fragment, directly generated from the bacmid via PCR or derived from sub-cloned transfer vectors if the fragment can be cut by seamless nuclease digest. Using the SwaI half sites analyzed above, the mutagenesis of the rAd genomes would consist of two steps. Firstly, the two SwaI half sites adjacent to the site of the intended mutation are complemented by red recombination and selected via antibiotic resistance testing introduced alongside completed half-sites, and secondly the intended sequence alongside the lost wt sequences are inserted (see Figure 1 A). As existing genomic features directly determine insertion sites, we decided to coin our methodology Half-site-directed Fragment Replacement (HFR).

To illustrate the general applicability of the system for adenoviruses, we chose to further analyze representative types of all human adenovirus species for the presence of half-sites of non-cutting enzymes. All analyzed viral genomes were found to be suitable for our approach, with an average insertion size ranging from 113 to 561 bp and potential maximum insertion sizes being from 1 kb to 4 kb, still permitting easy insertion of synthetic or PCR-derived DNA inserts (see supplementary data S4).

### Using HFR as a tool to generate single HAdV-C5 constructs expressing ZsGreen by tagging viral structural protein pIX

To demonstrate the utility of our system, we decided to generate recombinant viruses with a modified capsid protein pIX that is C-terminally fused a T2A-peptide (33) linked ZsGreen (34). To achieve this, we first replaced the genomic sequences between a forward SwaI half-site found in the E1B region and a backward half-site found in the inter-genic region between the pIX and the IVa2 genes by a Kanamycin-resistance-cassette (KnR) flanked by complemented SwaI sites (Figure 2 A). This resulted in an intermediate clone which can be linearized by SwaI digestion generating a linear genomic vector lacking the KnR fragment and harbors the pre-selected SwaI half-sites at each end (Figure 2 B). Overlapping insert fragments compatible with Gibson assembly were generated by PCRs using both viral DNA and the synthetic ZsGreen gene as a template (Figure 2 B). Notably, such overlapping sequences can be either found in the vector or added by primer design. After generating the intermediate pBWH-C5-pIX-Kan bacmid by red recombineering (16), it was SwaI digested and Gibson assembled with the two PCR derived fragments (Figure 2 C).

**Figure 2:**
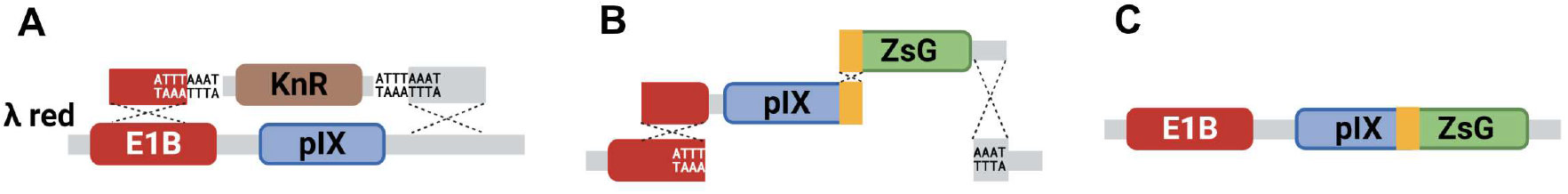
Schematic genetic workflow used for tagging pIX using a T2A peptide linker with ZsGreen; (A) replacement of pIX HFR locus with KnR; (B) reinsertion of pIX using PCRs fragments for both pIX and ZsGreen, connected via T2A (orange); (C) Finished modified pIX-locus with T2A-linked ZsGreen

This resulted in highly efficient assembly of recombinant bacmid, verified by both RFLP and Sanger sequencing. After rescue, four viral clones were selected and expanded. Extraction of viral DNA and subsequent RFLP analysis showed a length-shift from 3.2 kb to 4 kb, indicating the insertion of the transgene in all four single clones (Figure 3 A). We then verified ZsGreen expression after infection of 293a cells with a representative viral clone by fluorescent microscopy (Figure 3 B).

**Figure 3:**
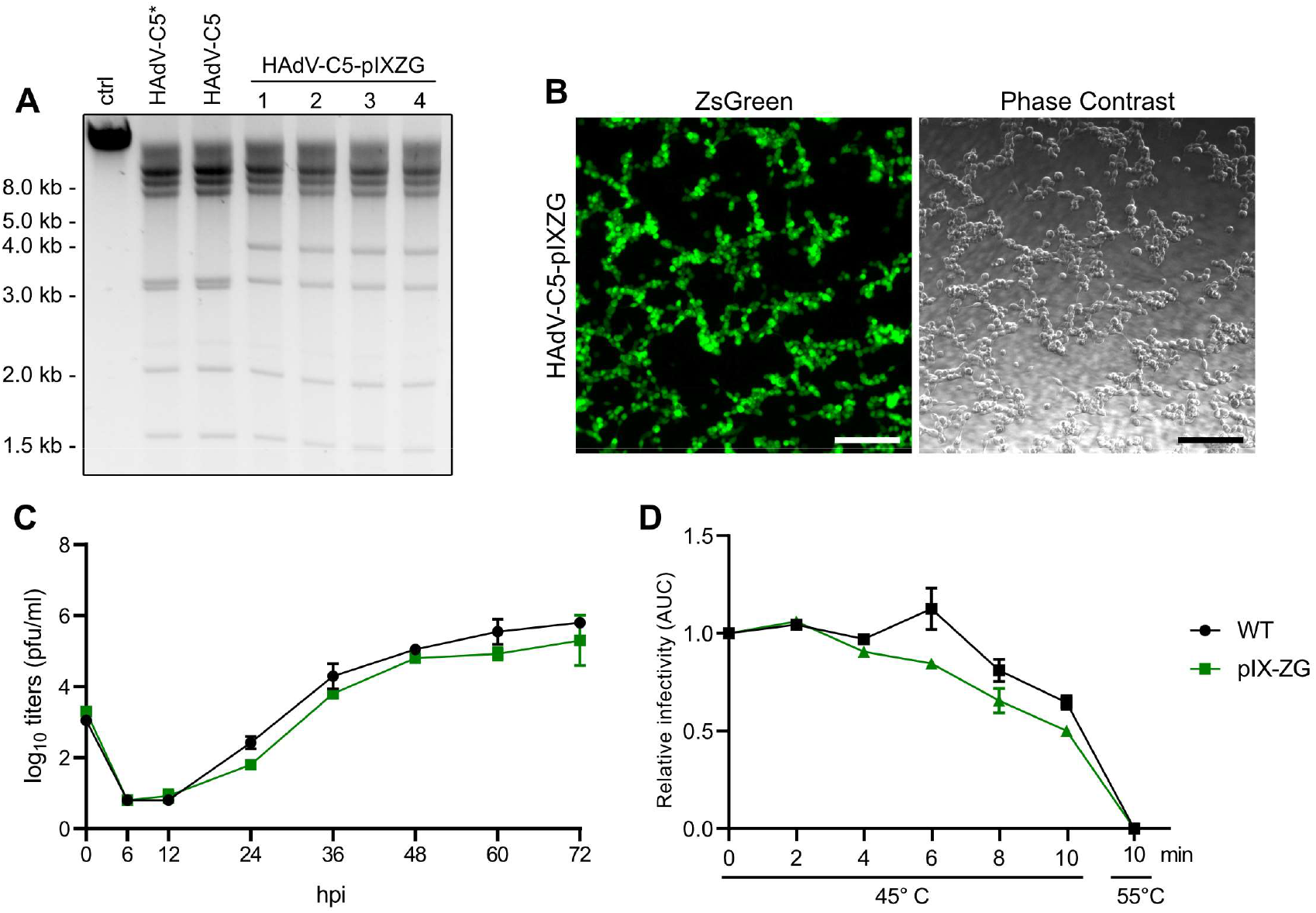
(A) RFLP analysis of extracted viral DNA, digested using DraIII. ctrl: undigested HAdV-C5 DNA derived directly from inoculum, HAdV-C5*: wildtype HAdV-C5 DNA derived directly from inoculum, HAdV-C5: wildtype-like HAdV-C5 DNA generated using CTR, HAdV-C5-pIXZG: recombinant HAdV-C5 DNA carrying ZsGreen-tagged pIX protein; (B) Micropscopy of HAdV-C5-pIXZG infected 293A cells using fluorescent channel and phase contrast. The scale bar represents 200 µm; (C) Growth curve of recombinant human Adenoviruses, comparing HAdV-C5 and HAdV-C5-pIXZG; (D) Thermostability experiment of recombinant human Adenoviruses, comparing HAdV-C5 and HAdV-C5-pIXZG

To analyze if our genetic modification affected viral growth or thermostability, growth kinetics and thermostability assays were performed using a representative viral clone in comparison to the unmodified WT virus. The pIX-ZG derived virus showed no significant growth attenuation or alteration in thermostability indicating that the genetically modified virus had no apparent fitness disadvantage in cell culture (Figure 3 C and D).

### Using HFR to generate libraries of barcoded rAd allows high recovery rates of possible assemblies at high efficiency

Next, we investigated the applicability of our workflow for the generation of viral genomic libraries. Instead of inserting a large library of transgenes or viral mutations, we decided to barcode a single site to assess the library’s diversity unbiased by downstream processes and viral fitness.

For the barcoding strategy, we wanted to achieve maximal efficiency and a complex randomization. Since the efficiency of Gibson assembly inversely correlates with the number of the fragments assembled and the size of the final construct, a single DNA fragment should contain the barcoded region. This ensures that the diversity is not affected by the number of inserts required during cloning. An additional advantage is that a single fragment derived from a small circular vector is more efficient and reliably cloned and thereby diversified compared to a bacmid derived DNA (36–38). Therefore, we designed a shuttle plasmid to release the artificial library in a single fragment. The pO6-fC5-GFP-MCS carried a DNA fragment consisting of the adenoviral packaging signal, a GFP transgene harboring a multiple cloning site (MCS) for introduction of the barcodes and part of the pIX locus (Figure 4 A). The insert was flanked by SapI sites to ensure seamless digestion.

**Figure 4:**
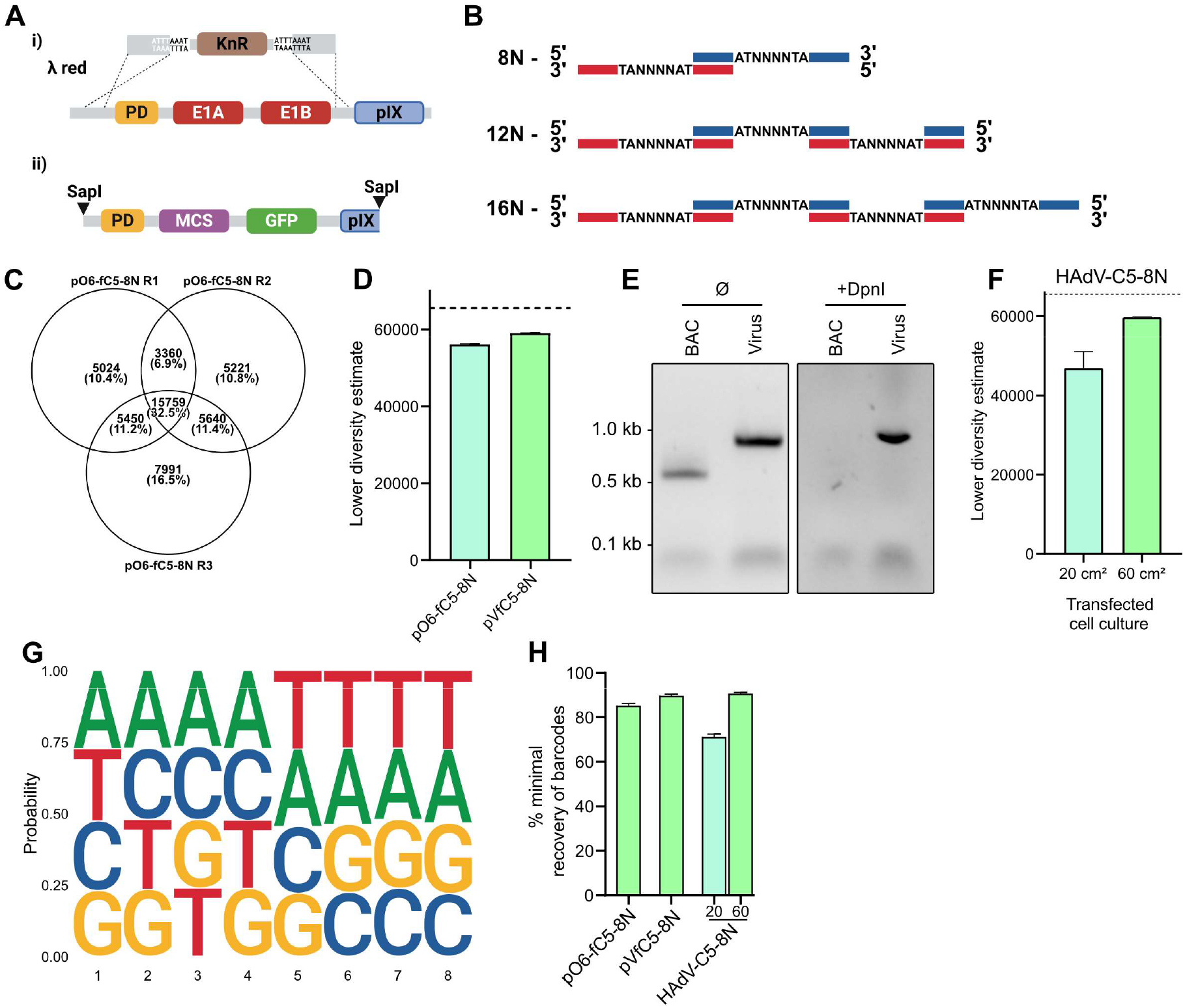
(A) Schematic of i) targeted region for transgenic insertion in bacmid, showing target removed region resulting in pBWH-fC5-E1Kn, ii) fragment released from pO6-fC5-GFP-MCS via SapI digestion, designated for bacmid insertion; (B) Overview of oligonucleotide-assemblies used for generation of diversified libraries, containing barcodes of 8, 12 or 16 nucleotides in length; (C) Schematic view of sequencing overlaps of barcodes recovered via NGS workflow, shown on three sequencing runs of pO6-fC5-8N; (D) Lower diversity estimate of plasmid (pO6-fC5-8N) and bacmid (pVfC5-8N) containing 8N-nucleotide diversified barcodes, the dashed line indiciates the maximum possible diversity; (E) Agarose gel showing PCR of extracted encapsidated DNA, using primers targeting either specifically bacmid or viral genomic DNA, pre- and post-DpnI digestion; (F) Lower diversity estimate of viral libraries containing 8N-nucleotide diversified barcodes, rescued in either 20 cm^2^ or 60 cm^2^ of transfected cell culture, the dashed line indicates the maximum possible diversity; (G) Frequency plot showing nucleotide frequency per position in barcode based on barcodes recovered from viral libraries. Sequences were aligned to strands shown above in Figure 4 B; (H) Overview of minimal estimated recovery rates for 8N-nucleotide based barcodes for different steps in the process, presented as percent of maximal possible diversity.

Next, we used overlapping oligonucleotides to assemble randomized barcodes and insert them into pO6-fC5-GFP-MCS (Figure 4 B). This was achieved by assembling two, three or four oligonucleotides each carrying a barcode consisting of 4 randomized nucleotides (8N, 12N and 16N). Using the 8N barcode gives a theoretical library diversity of 65,536. According to our previously published data (13) this diversity should not be limited by the virus rescue within smaller rescue experiments. To compare the theoretical diversity with that of the HFR generated libraries, we first generated a 8N-barcoded shuttle plasmid and transferred the insert into our bacmid. We sequenced each step in the library propagation process by next-generation sequencing (NGS) and analyzed the diversity with capture-recapture population modelling for heterogeneous populations as described by Chao (23), giving us an estimate for minimal diversity predicted within the analyzed sample (Figure 4 C). Sequencing was carried out on 200 bp PCR products of the barcoded region. Both pO6-subcloning and assembled bacmid-libraries showed similar levels of minimal expected barcode diversity at 5.61×10^4^ and 5.90×10^4^ unique barcodes, respectively (see Figure 4 D). The diversity was close to the theoretically achievable number of barcodes indicating that the diversity was likely not limited by the assembly workflow. Next, we rescued recombinant virus carrying the barcoded transgenes in either 20 cm^2^ or 60 cm^2^ cell culture dishes. Since our viral DNA samples were contaminated with a low level of bacmid-derived DNA even after benzonase treatment, we further purified our samples by DpnI digestion (Figure 4 E). As the DpnI recognition sequence is present in the PCR amplificate used for NGS, it should only affect non-encapsidated, bacmid-derived sequences due to the dam-methylated-GATC-specificity of the enzyme. Indeed, this treatment completely abolished PCR detection of bacmid- but not virus-derived DNA (Figure 4 E). The library diversity in the virus population was again assessed by NGS. The estimated diversity was with 4.69×10^4^ and 5.97×10^4^ unique barcodes for the 20 cm^2^ and 60 cm^2^ rescues highly similar to that of the pO6-subcloning and bacmid libraries (Figure 4 F). Nucleotide frequencies of the viral-encoded barcodes were randomly distributed indicating that libraries were not biased to specific barcodes (Figure 4 G). In each step of the library generation workflow, barcodes were consistently detected and reached a near-to-theoretical diversity. (Figure 4 H).

Therefore, we analyzed the limits of HFR by increasing the barcodes from 8N to 12N or 16N, (Figure 4 B), theoretically leading to diversities of 1.7×10^7^ and 4.3×10^9^ unique barcodes, respectively. Intriguingly, for both barcoding strategies we estimated between 10^6^ and 10^7^ unique barcodes for the pO6-subcloning and the bacmid libraries (Figure 5 A). As the library workflow reaches comparable levels of diversity in both bacmids, it is reasonable to assume that the bacmid libraries act as a limiting factor. This was substantiated by the recombinant virus rescue. 12N and 16N virus libraries produced from 20 cm^2^ or 60 cm² dishes resulted in diversity estimates of 5.17×10^5^/4.33×10^5^ and 1.44×10^6^/1.36×10^6^, respectively (Figure 5 B). Unlike 8N-coding libraries, these higher diversity libraries showed a clearer relation between surface area of transfected cell culture and barcode diversity. Notably, libraries from these transfections showed a position-independent bias towards thymine or the reverse complement adenine which likely represents a slight bias of the PCR or the synthesized nucleotides rather than a bias by the workflow itself (Figure 4 B, Figure 5 C,D). As both library preparations reach similar diversity estimates, despite having vastly different potential diversities, we can normalize these estimates by area of transfected cell culture used for generation libraries. This reveals a clear upper boundary for the here presented barcoding strategy of ∼2.5×10^4^ unique barcodes per cm^2^ of transfected cell culture (Figure 5 E). In summary, HFR combined with CTR produced highly diverse transgene libraries that were consistent over all steps in the assembly and rescue workflow.

**Figure 5:**
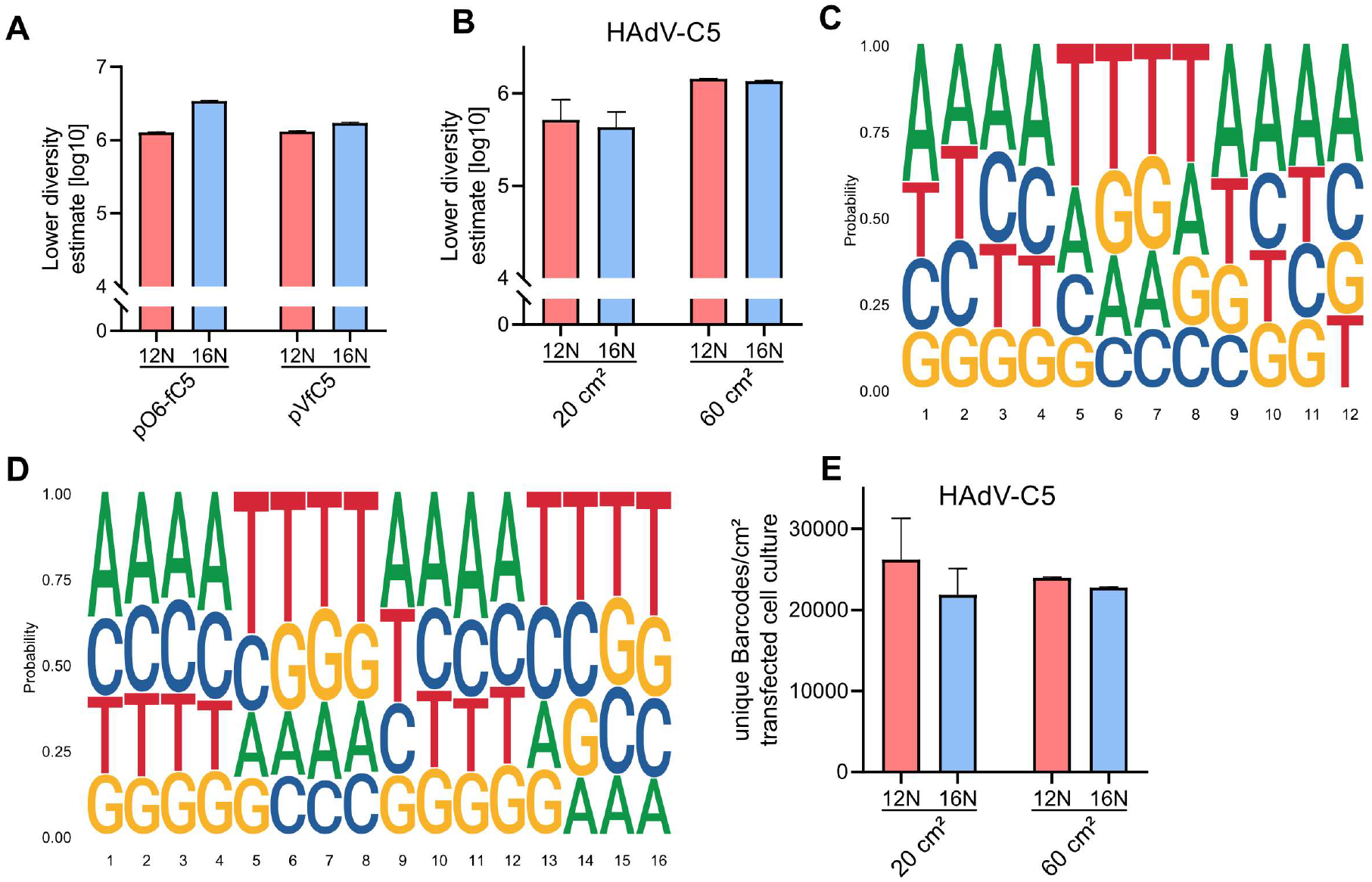
(A) Lower diversity estimates of pO6-based and bacmid cloning steps for libraries containing 12N- and 16N-diversified barcoded libraries; (B) Lower diversity estimate of viral libraries containing 12N- and 16N-nucleotide diversified barcodes, rescued in either 20 cm^2^ or 60 cm^2^ of transfected cell culture; (C) Frequency plot showing nucleotide frequency per position in barcodes based on barcodes recovered from 12N-diversified viral libraries; (D) Frequency plot showing nucleotide frequency per position in barcode based on barcodes recovered from 16N-diversified viral libraries; (E) Efficiency of barcode recovery for 12N- and 16N-diversified viral libraries normalized to cm^2^ of transfected cell culture.

### HFR modification and CTR rescue provide highly reliable viral rescues while preserving DNA integrity

One major concern during the generation of libraries is the presence of wildtype sequences that might be preferably selected compared to tailored particles. Using HFR as tool for generating libraries has the benefit of generating intermediate constructs, removing the possibility of contaminations with WT viruses. Attempting CTR with the intermediate pBWH-fC5-E1Kn construct yielded no viral particles due to removal of the essential packaging domain during the construction of the assembly vector. Using a clonal intermediate plasmid allows for efficient screening of successful selection of correct clones, removing potential background from recombineering. Utilizing transfer plasmids allows greater freedom in design, allowing addition of additional features such as restriction sites to ensure purity of plasmid library, removing background of miss-assembled libraries as well. Indeed, sequencing of viral DNA obtained from our library preparations against original plasmid revealed complete lack of WT sequences within mapped reads (see supplementary data S5). Furthermore, we also tested the overall genome integrity of the viral libraries to analyze for the presence of off-target-mutations. We were only able to detect mutations already present in the initial WT stock compared to the HAdV-C5-delE3 reference genome. Furthermore, frequencies of the detected mutations were highly similar in all samples indicating that the barcoded and WT rescued viruses were near-to-identical (Figure 6). A detailed overview of sequenced polymorphisms and their effect on CDS can be found in supplementary data S6. Notably, variants not occurring at 100% allele frequency were mostly called in highly repetitive genomic regions, which are notoriously difficult to amplify and sequence using established methods (39–41). In combination with the data obtained from single mutants analyzed for pIX-targeted mutagenesis, HFR did not cause off-target effects during library generation.

**Figure 6:**
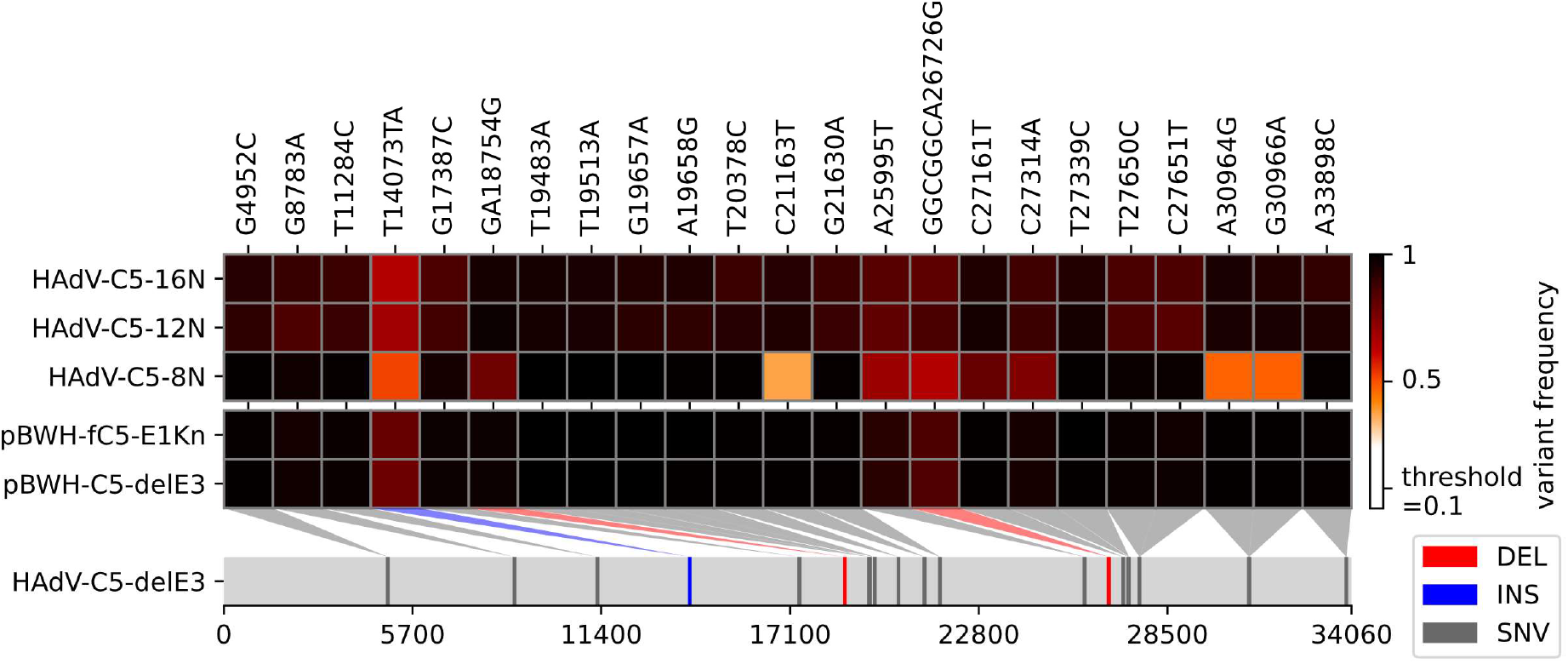
Variant frequency plot of analysed viral libraries showing variant frequency for detected SNPs and indels relative to HAdV-C5-delE3 viral reference genome.

## DISCUSSION

Handling and modifying large DNA genomes is not a trivial task. BAC-technology substantially aided this approach for large DNA viruses (11, 42–45). During recent years, a number of advances were made in this field by further improving the usability and flexibility of the bacmid-system (46–49). These innovations also spread to the field of rAds, often utilizing the high efficiency of Gibson assembly for streamlined adenoviral modification (31, 50–52). The major advantages of these new approaches are increased speed and reliability. In this work, we now further push the rAd technology by combing recent advancements with the flexibility of red recombination into a method coined HFR. We then apply this method in combination with our previously established CTR (13) to construct highly diverse rAd transgene libraries.

While previous work investigated the applicability of libraries in the adenoviral context, the workflow presented here improves both complexity of libraries by 10-100fold and the purity of generated rAd libraries as the whole viral population was to correctly recombined while previous work focused mostly on either high efficiency or high purity (53–55). A key feature of this novel technology is the flexibility in the modification site while minimizing off-target effects such as scars within the viral genome. Moreover, all steps of the library generation can be performed using commercially available standard reagents, also permitting a wide range of applications, as inexpensive oligonucleotides can be used for both diversification and linkage of individual fragments via PCR overlaps.

Using different lengths of randomized barcodes, the theoretical diversity varies by several orders of magnitude. Interestingly, we reached very similar levels of diversity in both higher-variance approaches. Therefore, we hypothesize that we have reached the maximal possible diversity of ∼7.5×10^4^ unique clones per ng of transformed subcloning vector DNA and ∼1.9×10^4^ unique clones per ng of bacmid DNA in our experimental setting (Figure 5). Since we used multiple fragments for the assembly of our shuttle plasmid, which is a less efficient approach compared to a two fragment assembly (56), the efficiency might be improvable. During cloning of single recombinant bacmids we found the primary determinant of cloning success being the nature of linear DNA fragment used for bacmid re-assembly (Figure 1 A/vi). Using enzymatically released fragments provided the most success here, so we suggest using the here shown method of generating specific sub-cloned libraries first. The bacmid assemblies on the other hand are already optimized, but linear upscaling could still increase diversity. However, great care must be taken when applying this method, as larger or smaller inserted cassettes might also impact efficiency.

Based on our prior work the observed reduced diversity after CTR compared to the prior assembly was expectable as rescue of rAd virus remains the major limiting factor (13). However, the here reached diversities are comparable to concurrent viral vector based library platforms utilizing lentivirus or AAV based vectors (57, 58) and are linearly scalable. In contrast to lentiviruses or AAVs, first generation adenoviruses are replication competent in the presence of E1 trans-complementation. Therefore, a linear scalability allows to increase library diversity similar to the expansion-based approaches used to overcome the limitations of bacteriophage vectors (59).

Using the lower-variability 8N approach for assembly, we were also able to recover a vast majority of available barcodes. We included this setting as proof-of-concept that HFR is usable for full-scale recovery of genomic libraries as long as total diversity of input is suitable for assembly and to show that the methodology to estimate the diversity was sound. The here used statistical capture/recapture model is an estimate of a total population size and typically under-rather than over-estimates population sizes (23). As such, actual recovery rate might be even higher than anticipated. However, manual inspection of the primary data obtained from the sequencing results seem to collaborate the obtained estimates.

Here, we used a transgene barcoding strategy which has no direct applicability as a proof-of-principle. Nevertheless, this technology can potentially be used for various library generation approaches such as CRISPR-forward screens or functional immunoglobulin screening. Even at the lowest estimate, we are able to achieve a comparable diversity to currently used AAV-technology (10). Using the same principles of feature-oriented mutagenesis utilized for AAV, we predict that rAds can be used in a similar fashion to further improve vector performance. HFR might be directly applicable for in vitro evolution based approaches of adenovirus capsids and might be a feasible alternative to currently available indirect applications based on phage displays (60, 61). However, reaching the levels of diversified functional virus shown here in in vitro evolution is unlikely as potentially vast amounts of variants might prove non-functional or display notably reduced fitness. Rational design still needs to be applied to library generation to ensure maximizing the possibility of selecting desired virus variants.

Other high-diversity applications, such as phage-display could also be tailored into rAd-libraries, allowing eukaryotic expression of target protein (62–64). This would improve both speed and applicability of the screening platform to allow faster advances in both research and medical purposes. Lentiviral libraries are already used in similar applications (65), however rAd provides a broad tropism, allowing for wide phenotypic screenings and monogenomic transductions (66, 67).

## DATA AVAILABILITY STATEMENT

The data underlying this article are available in the article and its online supplementary material. R code used for analysis is available online at https://doi.org/10.5281/zenodo.10040806. Raw data underlying these analyses is available online at https://doi.org/10.5281/zenodo.10090475.

## SUPPLEMENTARY DATA STATEMENT

Supplementary Data are available online.

## ACKNOWLEDGEMENT

We thank Harald Wodrich (Microbiologie Fondamentale et Pathogénicité, Bordeaux) for providing us with HH-Ad5-VI-wt and Albert Heim (Medizinische Hochschule Hannover, Institute of Biology, Hannover) for the wild type HAdV-C5. Hans-Gerhard Burgert for critical reading of the manuscript and his great mentorship along the way. We also want to acknowledge Simone Gruber and Dominique Gütle for providing excellent technical assistance.

