## supplementary data for "Combining CRISPR/Cas mediated terminal resolution with a novel genetic workflow to achieve high diversity adenoviral libraries"

### S1: Oligonucleotides used for diversification

Supplementary Data S1: Oligonucleotides used for generation of barcoding sequences. Diverse central regions are underlined. For 12N-library, oligonucleotide “Lib3_cap” was added to the mix to ensure that homologous regions would not be digested by 5’-Exonuclease.

| **Name** | **Sequence (5’ -> 3’)** |
|---|---|
| **Lib_Frag1** | ATATGTTTTATGTATCCAGTAACCATTGTANNNNATGACTCTAGAGGATCCCCGGGTACCGAGCTC |
| **Lib_Frag2** | TACAATGGTTACTGGATACATAAAACATATNNNNTAGTCCATCGGTTGCCCAAAGTGTTAAGAT |
| **Lib_Frag3** | ATTGCGGGAAACGGCCCTAGGGGTGATATANNNNATCTTAACACTTTGGGCAACCGATGGACTA |
| **Lib_Frag2_term** | TACAATGGTTACTGGATACATAAAACATATNNNNTAGACCTGCAGGCATGCAAGCTTGGCGTAATC |
| **Lib_Frag3_term** | GATTACGCCAAGCTTGCATGCCTGCAGGTCTANNNNATCTTAACACTTTGGGCAACCGATGGACTA |
| **Lib_Frag4_term** | TATATCACCCCTAGGGCCGTTTCCCGCAATNNNNTAGACCTGCAGGCATGCAAGCTTGGCGTAATC |
| **Lib3_Cap** | GACCTGCAGGCATGCAAGCTTGGCGTAATC |

### S2: Primers

Supplementary Data S2: Primers used for cloning in this study.

| Name | Sequence (5' - 3') |
|---|---|
| **BWHC05for** | TATTGGCTTCAATCCAAAATAAGGTATATTATTGATGATGCCTCCGGGGTCCACTGCAATTACTTCTCGACCAATTCTCATGTTTGAC |
| **BWHC05rev** | TATTGGCTTCAATCCAAAATAAGGTATATTATTGATGATGCCTCCGGGGTCCACTGCAATTATAAACTCGACAGCGACACACTTGC |
| **C5-pIXKan_for** | TTTGGGTAACAGGAGGGGGGTGTTCCTACCTTACCAATGCAATTTAAATTCGTGTGGGCGGACAATAAAGTCTTAAACTGAA |
| **C5-pIXKan_rev** | GCAAGACACTTGCTTGATCCAAATCCAAACAGAGTCTGGTTTTTTATTTAAATTGTGGGCGGACAAAATAGTTGG |
| **fC5-Kan-L_for** | GAATAAGAGGAAGTGAAATCTGAATAATTTAAATTCGTGTGGGCGGACAATAAAGTCTTAAACTGAA |
| **fC5-Kan-L_rev** | TTCCCACCCTTAAGCCACGCCCACACATTTAAATAAATGTGGGCGGACAAAATAGTTGG |
| **GHBfor** | CCGCGTGTGTACCTCTACCTGGAGTTTTTCCCACGGTGGA |
| **GHBrev** | TCCACCGTGGGAAAAACTCCAGGTAGAGGTACACACGCGG |
| **M13A_for** | GACGGGTAAAACGACGGCCAGT |
| **M13A_rev** | TAATGACTCAGTACAGGAAACAGCTATGAC |
| **O6-AVT_for** | AATGGAAGAGCTCCCATGTCAGCCGTTAAGTGTTCCTG |
| **O6-AVT_rev** | CATTGAAGAGCTTAGAAAAACTCATCGAGCATCAAATGAAACTGCAA |
| **O6-fC5_for** | GCTCGATGAGTTTTTCTAAGCTCTTCAATGGAATAAGAGGAAGTGAAATCTGAATAATTTGACGGGTAAAACGACGGCCAGT |
| **O6-fC5_rev** | CTTAACGGCTGACATGGGAGCTCTTCCATTTTCCCACCCTTAAGCCACGCCCACACATTTCAGTACCATAGAGCCCACCGCATCCC |
| **PCR-LibBB_for** | CATGCAAGCTTGGCGTAATCATGGTCA |
| **PCR-LibBB_rev** | GATCCCCGGGTACCGAGCTC |
| **pIXZsG_pIX_H3** | CGTCACCGCATGTGAGCAGACTTCCTCTGCCCTCTCCGGAAACCGCATTGGGAGGGGAGGAAGCCT |
| **pIXZsG_pIX_H5** | CTACCTTACCAATGCAATTTGAGTCACACTAAGATATTGCT |
| **pIXZsG_ZG_H3** | CAGAGTCTGGTTTTTTATTTATGTTTCAGGGCAAGGCGGAGCCGGAG |
| **pIXZsG_ZG_H5** | TCTGCTCACATGCGGTGACGTGGAGGAGAATCCCGGGCCAGCCCAGTCCAAGCACGGCCTGAC |

**S3: Plasmids**

Supplementary Data S3: Plasmids used and generated during this study, including size and notably features.

| Plasmid Name | Size | Features | Accession |
|---|---|---|---|
| pGPS1.1 | 4814 bp | GPS-1 Genome Priming System transposon donor; kanamycin resistance (KanR); tetracycline resistance (TetR) | 10666445 |
| pO6-A5-GFP | 3457 bp | Constitutive GFP expression under human CMV promoter; HAdV-C5-targeted transfer vector, carrying part of packaging domain; kanamycin resistance (KanR) (1) | |
| pKD46 | 6329 bp | red-recombineering plasmid; Arabia sugar dependent expression of lambda red phage exo, beta and gam protein; temperature sensitive; β-lactamase expression (AmpR) | 10829079 |
| pO6-fC5-GFP | 3047 bp | Constitutive GFP expression under murine CMV promoter; HAdV-C5-targeted transfer vector, carrying part of packaging domain; empty multi cloning site; kanamycin resistance (KanR) | OR810920 |
| pO6-fC5-8N-GFP | 3089 bp | Constitutive GFP expression under murine CMV promoter; HAdV-C5-targeted transfer vector, carrying part of packaging domain; multi cloning site equipped with 2-oligo diversified barcode; kanamycin resistance (KanR) | OR810926 |
| pO6-fC5-12N-GFP | 3123 bp | Constitutive GFP expression under murine CMV promoter; HAdV-C5-targeted transfer vector, carrying part of packaging domain; multi cloning site equipped with 3-oligo diversified barcode; kanamycin resistance (KanR) | OR810927 |
| pO6-fC5-16N-GFP | 3157 bp | Constitutive GFP expression under murine CMV promoter; HAdV-C5-targeted transfer vector, carrying part of packaging domain; multi cloning site equipped with 4-oligo diversified barcode; kanamycin resistance (KanR) | OR810928 |
| pBWH-C5-delE3 | 40498 bp | Genomic BACmid; carrying mutant HAdV-C5 genome (HH-Ad5-VI-wt(2)), E3-region deleted; ITRs flanked by ACT sequences; chloramphenicol resistance (CamR) | OR810922 |
| pBWH-fC5-E1Kn | 39014 bp | Genomic BACmid; carrying HAdV-C5 genome, E3-region deleted; E1-region replaced with KanR to facilitate HFR; ITRs flanked by ACT sequences; chloramphenicol resistance (CamR); kanamycin resistance (KanR) | OR810925 |
| pVfC5-8N | 38946 bp | Genomic BACmid; carrying HAdV-C5 genome, E1/E3-region deleted; carrying 2-oligo diversified genetic barcoding region; Expression of GFP under murine CMV promoter; ITRs flanked by ACT sequences; chloramphenicol resistance (CamR) | OR810929 |
| pVfC5-12N | 38980 bp | Genomic BACmid; carrying HAdV-C5 genome, E1/E3-region deleted; carrying 3-oligo diversified genetic barcoding region; Expression of GFP under murine CMV | OR810930 |

| | | | |
|---|---|---|---|
| | | promoter; ITRs flanked by ACT sequences; chloramphenicol resistance (CamR) | |
| pVfC5-16N | 39014 bp | Genomic BACmid; carrying HAdV-C5 genome, E1/E3-region deleted; carrying 4-oligo diversified genetic barcoding region; Expression of GFP under murine CMV promoter; ITRs flanked by ACT sequences; chloramphenicol resistance (CamR) | OR810931 |
| pAR-Int5-Cas9 | 6699 bp | Constitutive sgRNA expression under human U6 promoter, targeting 5' of HAdV-C5 ITR sequence; Constitutive SpCas9-FLAG-NLS expression under EF1α-promoter; ampicilin resistance | - |
| pBWH-C5-pIX-Kan | 43300 bp | Genomic BACmid; carrying HAdV-C5 genome; pIX locus replaced with KanR to facilitate HFR; ITRs flanked by ACT sequences; chloramphenicol resistance (CamR); kanamycin resistance (KanR) | OR810923 |
| pBWH-C5-pIXZG | 43216 bp | Genomic BACmid; carrying HAdV-C5 genome; pIX protein C-terminally linked to ZsGreen (3) via T2A self-cleaving peptide; ITRs flanked by ACT sequences; chloramphenicol resistance (CamR) | OR810924 |
| pBWH-C5 | 42376 bp | Genomic BACmid; carrying HAdV-C5 genome; ITRs flanked by ACT sequences; chloramphenicol resistance (CamR) | OR810921 |
| pKSB2 | 6457 bp | Single-copy BACmid backbone; chloramphenicol resistance (CamR) | - |

### S4: Overview of exemplary HAdV-Species and applicable enzymes for HFR

Supplementary Data S4: *In silico* analysis of representative genomes for each human adenovirus species for applicability of HFR mutagenesis.

| Virus | GenBank | Candidate Enzyme | Forward Sites | Min. Distance | Max. Distance | Mean | Reverse Sites | Min. Distance | Max. Distance | Mean | Minimum possible insertions | Min. Size | Max. Size | Mean |
|---|---|---|---|---|---|---|---|---|---|---|---|---|---|---|
| A12 | NC001460 | SnaBI (TACGTA) | 534 | 3 | 606 | **76,27** | 454 | 3 | 859 | **75** | 988 | 2 | 1004 | **113,49** |
| | | SwaI (ATTTAAAT) | 231 | 4 | 1267 | **147,24** | 224 | 5 | 1109 | **174,11** | 454 | 2 | 1361 | **231,26** |
| B11 | AF532578 | SwaI (ATTTAAAT) | 174 | 4 | 1096 | **199,38** | 188 | 5 | 1040 | **184,53** | 361 | 2 | 1476 | **296,8** |
| | | PmeI (GTTTAAAC) | 161 | 8 | 1632 | **213,99** | 189 | 4 | 1437 | **181,32** | 350 | 2 | 1906 | **345,75** |
| B3 | DQ086466 | SwaI (ATTTAAAT) | 144 | 5 | 1900 | **244,73** | 171 | 5 | 1657 | **206,09** | 315 | 3 | 2145 | **349,52** |
| | | PmeI (GTTTAAAC) | 128 | 9 | 1665 | **272,95** | 164 | 4 | 2081 | **212,95** | 291 | 2 | 2176 | **407,09** |
| C2 | AC000007 | SnaBI (TACGTA) | 427 | 3 | 823 | **83,89** | 357 | 3 | 1100 | **100,34** | 785 | 2 | 1104 | **165,99** |
| C5 | AC000008 | SwaI (ATTTAAAT) | 125 | 6 | 2093 | **286,1** | 136 | 5 | 2423 | **262,3** | 262 | 5 | 3191 | **430,23** |
| C5delE3 | | SwaI (ATTTAAAT) | 112 | 6 | 2093 | **302,54** | 125 | 5 | 2432 | **270,36** | 238 | 6 | 3191 | **447,95** |
| D17 | AF108105 | SwaI (ATTTAAAT) | 98 | 4 | 3242 | **257,06** | 122 | 4 | 2828 | **286,82** | 217 | 2 | 3369 | **470,04** |
| | | PmeI (GTTTAAAC) | 90 | 6 | 1852 | **389,08** | 133 | 8 | 2298 | **263,29** | 222 | 2 | 2592 | **536,46** |
| | | SnaBI (TACGTA) | 368 | 3 | 1022 | **95,27** | 308 | 3 | 848 | **113,82** | 677 | 2 | 1096 | **192,49** |
| E4 | AY458656 | SwaI (ATTTAAAT) | 100 | 4 | 2293 | **359,21** | 109 | 4 | 3778 | **329,55** | 210 | 3 | 3947 | **561,12** |
| F41 | DQ315364.2 | PmeI (GTTTAAAC) | 192 | 5 | 1748 | **176,98** | 218 | 5 | 959 | **155,89** | 409 | 3 | 2000 | **275,49** |
| G52 | DQ923122.2 | SwaI (ATTTAAAT) | 123 | 4 | 2017 | **375,46** | 98 | 4 | 3249 | **345,16** | 221 | 2 | 4003 | **471,08** |

**S5: Mapping and coverage of genome reads obtained from library preparations against original virus**

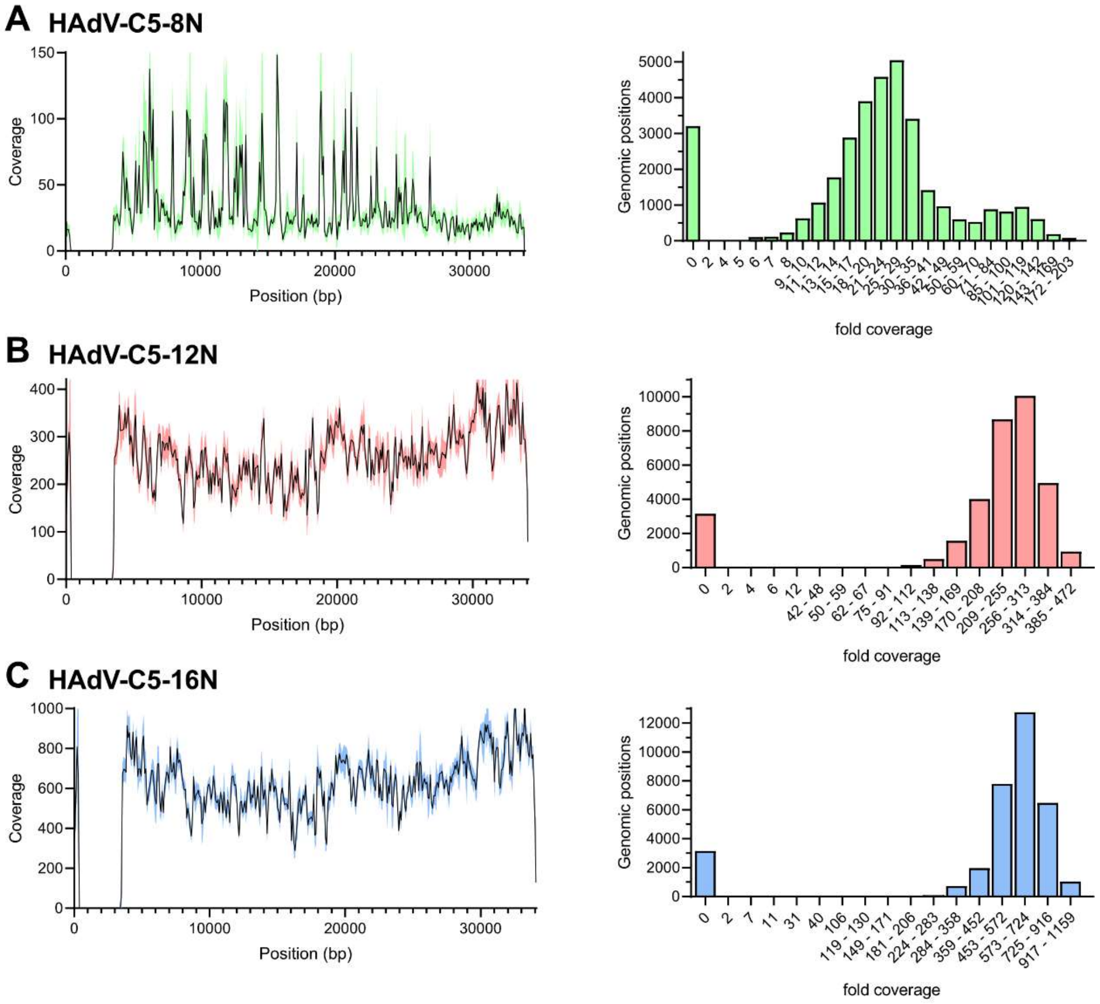


Supplementary Data S5: Mapping statistics for NGS reads obtained for viral genomic DNA directly after rescue. Data shows coverage of genetic region by mapping location (left) and nucleotide amount (right) for recombinant virus containing (A) 8N, (B) 12N- and (C) 16N-diversified viral libraries.

### S6: Overview of mutations relative to reference detected using NGS

Supplementary Data S6: Overview of variants detected during bacmid and rAd genome sequencing, relative to HAdV-C5-delE3 mutant virus reference sequence.

| Position | Mutation | CDS |
|---:|---|---|
| **4952** | SNP | pIVa2 |
| **8783** | Ala -> Asp | pTP/Pol |
| **11284** | Tyr -> His | 52K |
| **14073** | polyA | NCR |
| **17387** | Gly -> Arg | pV |
| **18754** | TGA*A -> TGA* | NCR |
| **19483** | SNP | Hexon |
| **19513** | SNP | Hexon |
| **19657** | SNP | Hexon |
| **19658** | Thr -> Ala | Hexon |
| **20378** | SNP | Hexon |
| **21163** | SNP | Hexon |
| **21630** | Arg -> Glu | Hexon |
| **25995** | SNP | 100K |
| **26726** | Pro/Ala/Ala -> Pro | 33K |
| | Gly/Gly/Ser -> Gly | 22K |
| **27161** | SNP | NCR |
| **27314** | SNP | pVIII |
| **27339** | Leu -> Pro | pVIII |
| **27650** | SNP | pVIII |
| **27651** | Pro -> Ser | pVIII |
| **30964** | SNP | NCR |
| **30966** | SNP | NCR |
| **33898** | SNP | NCR |
